# Rational molecular design for improved ZHD101 thermal stability based on the introduction of disulfide bonds at the dimer interface and B-factor analysis

**DOI:** 10.1101/2024.12.11.628026

**Authors:** Weiqiu Ding, Yujie Huang, Haiyi Zhang, Shaoyan Zheng, Chunfang Xie, Dongsheng Yao

## Abstract

Zearalenone hydrolase 101 (ZHD101), derived from *Clonostachys rosea*, is known to effectively degrade the main contaminant (zearalenone) in animal feed, but the thermal instability of ZHD101 limits its industrial application. In this study, we successfully enhanced the thermal stability of ZHD101 through two iterative rounds of rational molecular design. First, after the prediction of disulfide bond sites, ZHD101^T229C^ was obtained, and a new disulfide bond was formed between two single ZHD101 subunits to construct a dimer ZHD101. Then, based on ZHD101^T229C^, two high-vibration hotspot amino acids N137 and D170 were selected by analyzing atomic position fluctuations and dynamic information A small and precise mutant library containing three mutants (ZHD101^T229C/N137L^, ZHD101^T229C/D170L^, and ZHD101^T229C/D170C^) was obtained by saturation mutagenesis and calculation of binding energy in silico. Compared with the wild type, the thermal half-inactivation temperature (*T_50_*) of ZHD101^T229C/D170C^ increased by 7°C, its half-life (*t_1/2_*) increased by 200% at 50°C, and its melting temperature (*T_m_*) increased by 18.1°C. Molecular docking suggested that new covalent bond formation and shorter bond distance may contribute to the improved thermal stability of ZHD101. The rational design strategy proposed in this work can provide a reference for the thermal stability modification and optimization of other proteins.

**Importance:** Zearalenone (ZEN) is a nonsteroidal estrogenic mycotoxin that poses a significant threat to animal feed safety. ZEN hydrolase 101 (ZHD101) catalyzes the conversion of ZEN into a nontoxic product, offering a promising detoxification strategy. However, the thermal instability of ZHD101 severely limits its industrial application. In this study, the thermal stability of ZHD101 was significantly improved through two rounds of rational design, resulting in the ZHD101^T229C/D170C^ mutant. ZHD101^T229C/D170C^ exhibited the improved half-inactivation temperature (*T_50_*), half-life (*t_1/2_*) and the highest melting temperature (*T_m_*) reported thus far. Overall, the iterative combinatiorial mutation strategy involved the introduction of a disulfide bond between two single subunits and the application of B-factor analysis to identify hotspot amino acids residues. This approach effectively enhanced the enzyme’s structural stability, paving the way for its industrial application. ZHD101^T229C/D170C^ and the rational design strategy presented in this work provide a robust framework for the thermal stability optimization of similar biocatalysts, advancing the practical use of ZHD101 in the feed industry.

---

Zearalenone (ZEN) contamination poses a major threat to the global stockbreeding and food industries [1–4]; ZEN is a key contributor to animal feed pollution[5–8]. ZEN is a nonsteroidal estrogenic mycotoxin that adversely affects the reproductive system of female animals, causing infertility and slow growth [9, 10]. Furthermore, ZEN can accumulate and propagate through the food chain, ultimately posing health risks to humans [11, 12]. Enzymatic has emerged as a highly efficient and specific approach for reducing feed contamination by ZEN [13]. Zearalenone hydrolase (ZHD101, GenBank ID: AB076037.1), identified from the fungus *Clonostachys rosea*, was the first enzyme reported to degrade ZEN [14]. ZHD101 is considered the most effective enzyme for ZEN degradation due to its ability to break the lactone bond in ZEN’s structure. This structural disruption prevents ZEN from binding to estrogen receptors, thereby neutralizing its toxicity [15]. Moreover, ZHD101 has also been reported to be the only α/β-hydrolase that can detoxify ZEN and its derivatives[16]. However, the optimal temperature for ZHD101 is between 37°C and 45°C, and it rapidly loses activity at 50°C, showing poor thermal stability, which greatly limits its application as a feed additive [17–20]. Addressing this limitation has become a focal point for researchers seeking to enhance the industrial viability of ZHD101.

As the temperature increases, heat causes enhanced vibration and motion of protein conformations, ultimately affecting the enzyme function [21–23]. Therefore, the key to improving the thermal stability of enzymes is enhancing the rigidity of their molecular structures, especially their spatial structures while preserving catalytic function. Conventional strategies used to enhance structural stability include introducing disulfide bonds [24, 25], hydrogen bonds [26], salt bridges [27–29], and hydrophobic interactions [30] within the molecule to strengthen internal interactions; introducing proline to disrupt flexible structures such as helices or loops to increase protein rigidity [31]; and fusing the protein with a known thermostable protein domain to improve its stability [32].

Some researchers have used the abovementioned strategies to improve the thermal stability of ZHD101 and achieved positive results. However, these molecular modifications were performed on single ZHD101 subunit. Notably, many proteins function as polymers rather than single protein subunit, such as tetrameric lactate dehydrogenase and dimeric glutathione reductase. Studies have revealed that multimeric enzymes’ activity and stability are better than monomeric enzymes [33], and their quaternary structure may endow proteins with new properties. Peng et al. successfully analyzed the oligomerization status of ZHD101 in solution and its crystal structure, and observed an apparent molecular weight of 55.3 kDa, indicating that ZHD101 exists as a homodimer in solution [16].Therefore, the ZHD101 monomer may only function after being aggregated into a polymer through non-covalent bonds.

Based on the above considerations, this work aims to enhance the thermal stability of ZHD101 by introducing disulfide bond between two single ZHD101 subunits to form dimer and improving the rigidity of its quaternary structure. First, disulfide bonds were design at the subunit interface, facilitating the connection of ZHD101 monomers into a dimer with enhanced structural rigidity. Next, amino acid residues with unstable and high degrees of freedom in the dimer were identified using the B-factor analysis. A mutant screening library by virtual saturation mutagenesis of unstable amino acid. The double mutants were selected based on their lowest binding energy. Finally, the thermal stability of the double mutants was experimentally validated and characterized. We successfully obtained ZHD101 mutants with significantly improved thermal stability, demonstrating the feasibility and effectiveness of our rational design strategy for enzyme optimization. The evolved enzyme can better meet industrial production needs, especially in stockbreeding and food industries.

## Results and discussion

### Selection of inter-subunit disulfide bond formation sites

As a covalent bond, the bonding energy of disulfide bonds is much greater than that of other secondary bonds. The introduction of disulfide bonds can reduce the entropy of the protein skeleton and improve the rigidity of the protein structure[34, 35]. The introduction of disulfide bonds is not randomly designed, and the wrong position of introduction may reduce the structural stability.

The interface between the two subunits in a dimer is ideal for disulfide bond introduction. Results indicated that the interface of the ZHD101 dimer was composed of 14 amino acids, among which F222, I225, V226, I235, and L237 formed a stable hydrophobic surface, which helped stabilize the dimer (Fig 1). Molecular docking and alanine scanning results showed that S102, L135, W183, F221, and H242 were the key amino acids at the binding surface of ZHD101 and ZEN (Fig S1 and Table S1). The subsequent mutation design avoided the amino acid residues.

**FIG 1.**
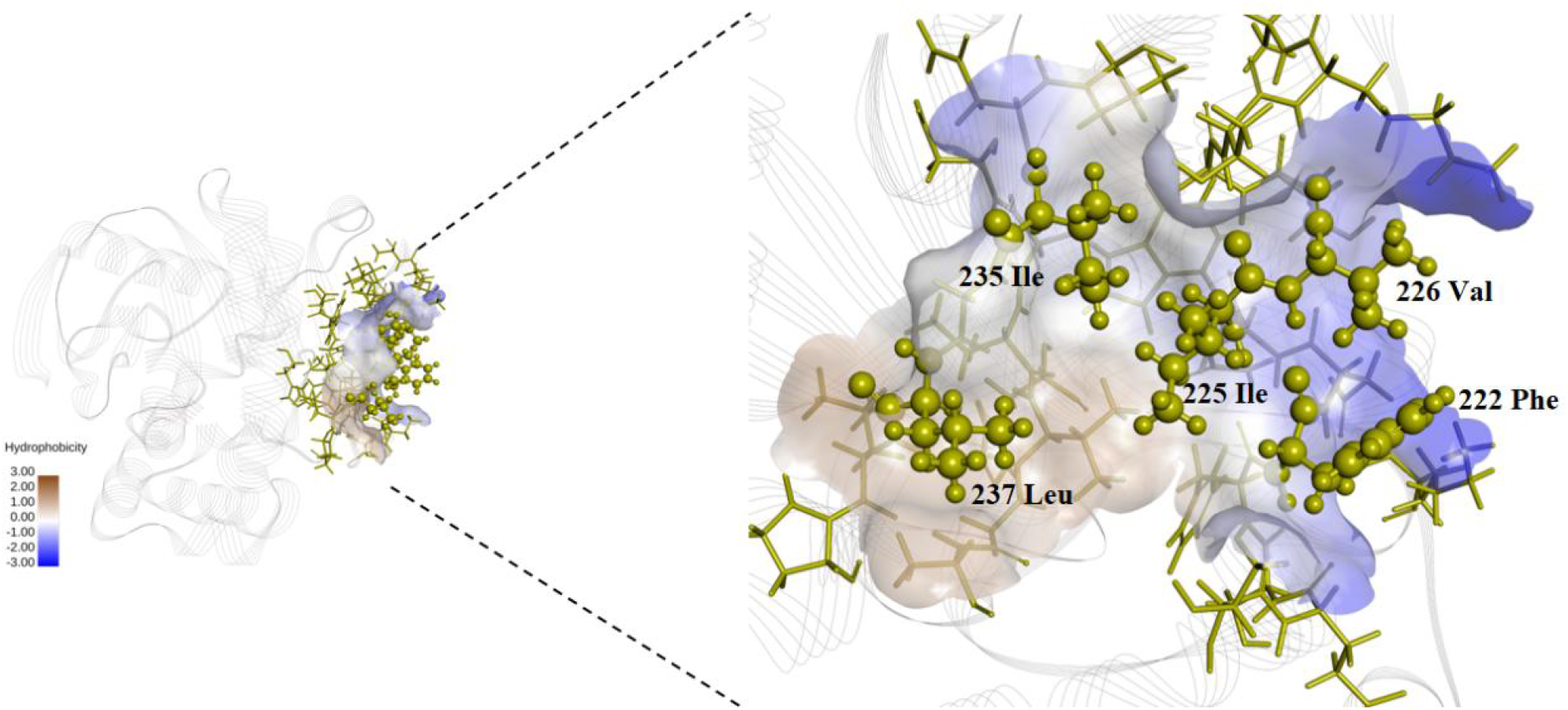
The hydrophobic surface on the dimer interface of ZHD101 is formed by F222, I225, V226, I235, and L237.

Using DS2023 for disulfide bond prediction (Fig 2), we observed that the mutant that may form interchain disulfide bonds was A: T229C/B: T229C (ZHD101^T229C^), indicating that the 229th threonine of both monomers (named A and B) was mutated to cysteine. The bond energy for the formation of interchain disulfide bonds was 5.38 kcal/mol, and the Chi (χ3) torsion angle was 100.6 °. The other mutant was A: T216C/B: T218C (ZHD101^T216C/T218C^), but its bond energy for forming interchain disulfide bonds was 9.96 kcal/mol, and the Chi(χ3) torsion angle was -127.9 °, indicating that disulfide bond formation is somewhat difficult or the resulting dimer may have poor stability (Table S2).

**FIG 2.**
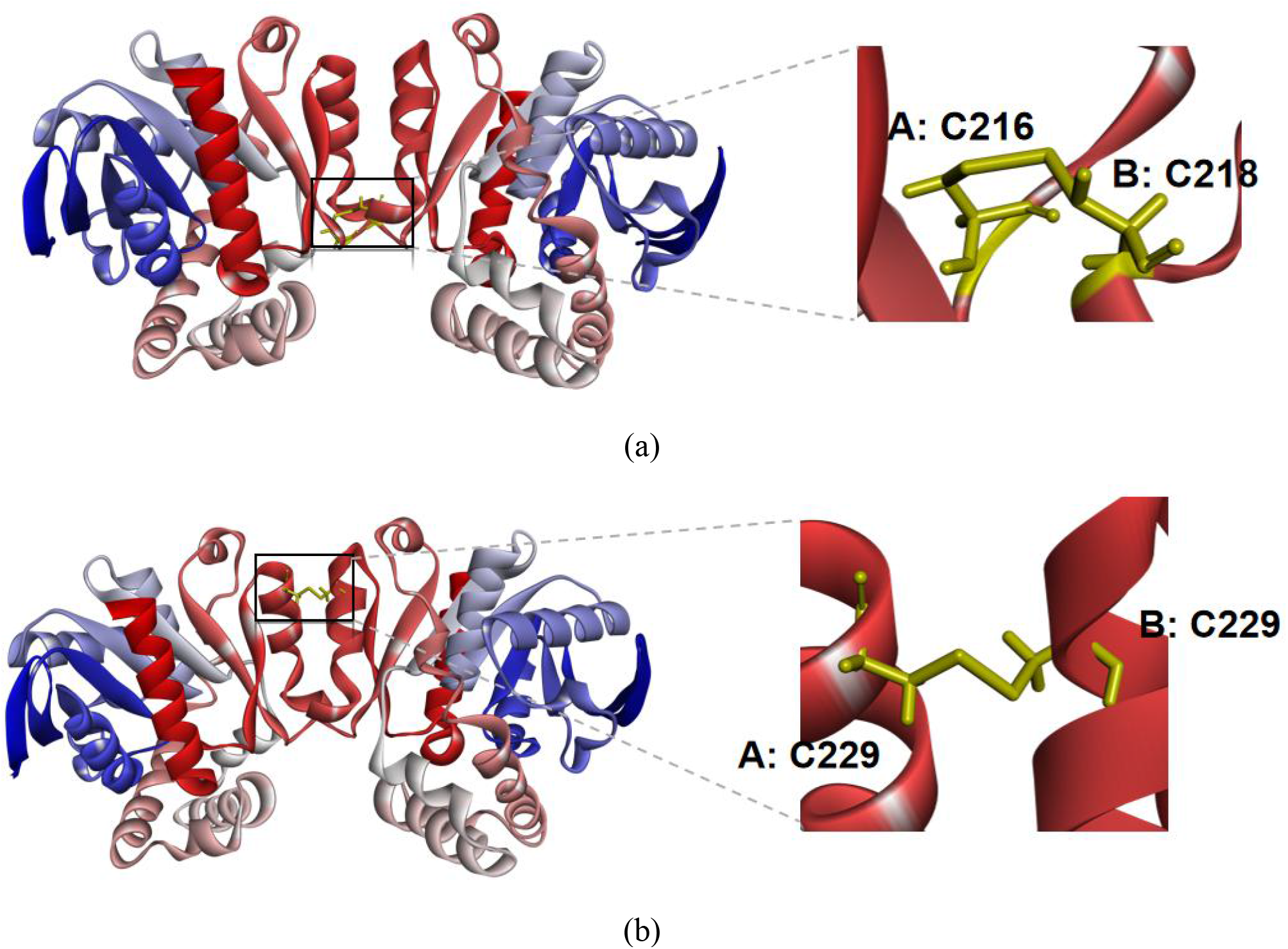
Disulfide bond formation site between subunits of mutant enzymes. (a) ZHD101^A216C/E218C^. (b) ZHD101^T229C^.

Using DbD2 for disulfide bond prediction, we found that A: T229C/B: T229C (ZHD101 ^T229C^) and A: A214C/B: E219C (ZHD101 ^A214C/E219C^) formed interchain disulfide bonds. The B-factor of the protein region containing the A: T229C/B: T229C candidate disulfide bond was 93.05, which was higher than that of A: A214C/B: E219C (Table S3). The mutation energies of mutants ZHD101^T229C^ (-0.49 kcal/mol), ZHD101^T216C/T218C^ (-0.38 kcal/mol), and ZHD101^A214C/E219C^ (-2.36 kcal/mol) were less than 0.5 kcal/mol, indicating a relatively small impact on the stability of enzyme molecular structure after mutation. Based on the above results, the mutants ZHD101^T229C^, ZHD101^T216C/T218C^, and ZHD101^A214C/E219C^ were selected for subsequent analysis of disulfide bond formation and thermal stability. Three methods (DS2023, DBD2, and B-factor) were used to predict the mutation site of the disulfide bond. DS2023 had the highest accuracy and could effectively improve the efficiency of screening mutants.

### Identification of inter-subunit disulfide bonds

Since disulfide bonds can be reduced. By adding β-mercaptoethanol in the loading buffer, SDS-PAGE was used to compare the molecular weights of reduced and non-reduced proteins to determine whether inter-monomer disulfide bonds were formed. ZHD101^T229C^ exhibited a band at approximately 55–60 kDa after electrophoresis without a reducing agent, which was twice the size of the monomer (28 kDa), demonstrating the successful formation of dimers through the introduction of disulfide bonds (Fig S2(a)).

In addition, we observed a 28 kDa band, indicating the presence of monomeric proteins, which may be those that have not yet formed dimers or may have been formed by the reduction of dimers. ZHD101^A214C/E219C^ did not successfully form disulfide bonds; however, the introduction of site-specific glycosylation promoted the formation of disulfide bonds (data not shown). ZHD101^T216C/T218C^ was not successfully expressed. We proceeded with the subsequent rational molecular design of the B-factor in ZHD101^T229C^.

### Prediction of mutation sites for improving thermostability based on B-factor

The B-factor, sometimes referred to as the temperature factor or atomic shift parameter, is used in protein crystallography to describe the attenuation of X-rays or neutron scattering caused by thermal motion. The B-factor can be used to identify the flexibility of atoms, side chains, or even entire regions in a protein.

The thermal vibration status of each amino acid in the dimer ZHD101^T229C^ was analyzed using the B-factor (Fig S3); the top 20 sites with the highest B-factor values and largest thermal vibration amplitudes are listed in Table S4. After virtual amino acid mutation, we selected 10 mutation sites with mutation energies less than -2 kcal/mol and lower energy (N137L, D199Q, D170L, A139R, D170C, E163Q, D199W, D143Q, D133W, D170W) for subsequent evaluation (Table S5). Using the DUET server to calculate the ΔΔG in folding free energy between the wild type and mutants, mutants with lower and more stable energy were screened: ZHD101^T229C/N137L^ (0.304 kcal/mol), ZHD101^T229C/D170L^ (0.501 kcal/mol), and ZHD101^T229C/D170C^ (0.216 kcal/mol). After experimental verification, these mutants formed intermolecular disulfide bonds and successfully constructed dimers (Fig S2(b)); next, we performed protein structure and thermal stability analyses.

### Thermal stability analysis

The methods described above studied the thermostabilities of the wild-type and mutant enzymes. As shown in Fig 3(a), after incubation at 20℃ for 10 min, the wild-type enzyme activity was considered 100%, and the other groups’ relative activity was calculated. The thermal half-inactivation temperature (*T_50_*) of ZHD101^T229C/D170C^ was approximately 54℃, 7℃ higher than that of the wild type. The *T_50_* of the other mutants was lower than that of the wild type or showed no significant difference. As shown in Fig 3(b), At 50℃, the half-life (*t_1/2_*) of ZHD101^T229C^ and ZHD101^T229C/D170C^ was 180% and 200% of that of the wild type, respectively. These results indicate that introducing disulfide bonds at the subunit interface to construct dimers, combined with a subsequent B-factor rational design, can significantly enhance the thermal stability of the enzyme.

**FIG 3.**
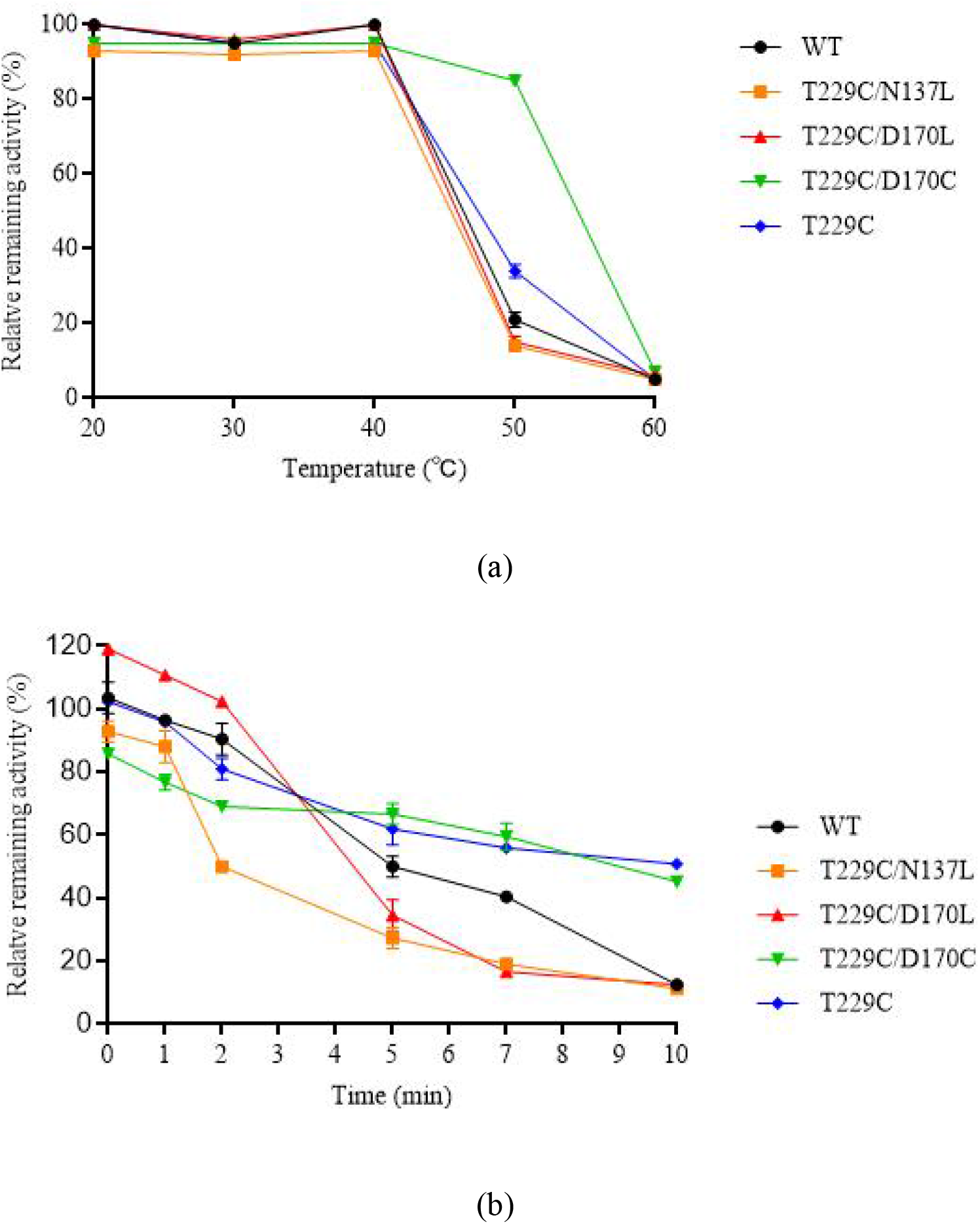
Thermal stability of ZHD101 and mutants. (a) *T50* of ZHD101^WT^, ZHD101^T229C/N137L^, ZHD101^T229C/D170L^, ZHD101^T229C/D170C^and ZHD101^T229C^ (b) *t1/2* of ZHD101^WT^, ZHD101^T229C/N137L^, _ZHD101_T229C/D170L _, ZHD101_T229C/D170C_and ZHD101_T229C_._

Additionally, ZHD101^T229C/D170L^ exhibited markedly higher initial enzyme activity than the wild type, but had lower thermal stability, making it a potential starting enzyme for further modification. The full wavelength circular dichroism spectra showed the protein folding of the mutant was consistent with that of the wild type (Fig 4). The results of thermal denaturation showed that the melting temperature (*T_m_*) of the wild type was 49℃ (Fig S4(a)), while ZHD101^T229C/D170C^ had two different *T_m_* values, *T_m1_* at 50.5℃ and *T_m2_* at 67.1℃ (Fig S4(b)). The two *T_m_* values were attributed to the ZHD101^T229C/D170C^ sample containing a mixture of dimers and monomers. *T_m1_*, which was close to the wild-type *T_m_*, corresponded to the monomeric form of ZHD101^T229C/D170C^, whereas *T_m2_* corresponded to the dimeric form of ZHD101^T229C/D170C^. Compared to the wild type, the *T_m_* of the dimer increased by 18.1℃ (*ΔT_m_*=18.1℃). The thermal stability of the dimer formed by ZHD101^T229C/D170C^ was significantly enhanced. In recent years, many researchers have used different methods to modify and characterize the thermal stability of ZEN, as summarized in Table 1.

**FIG 4.**
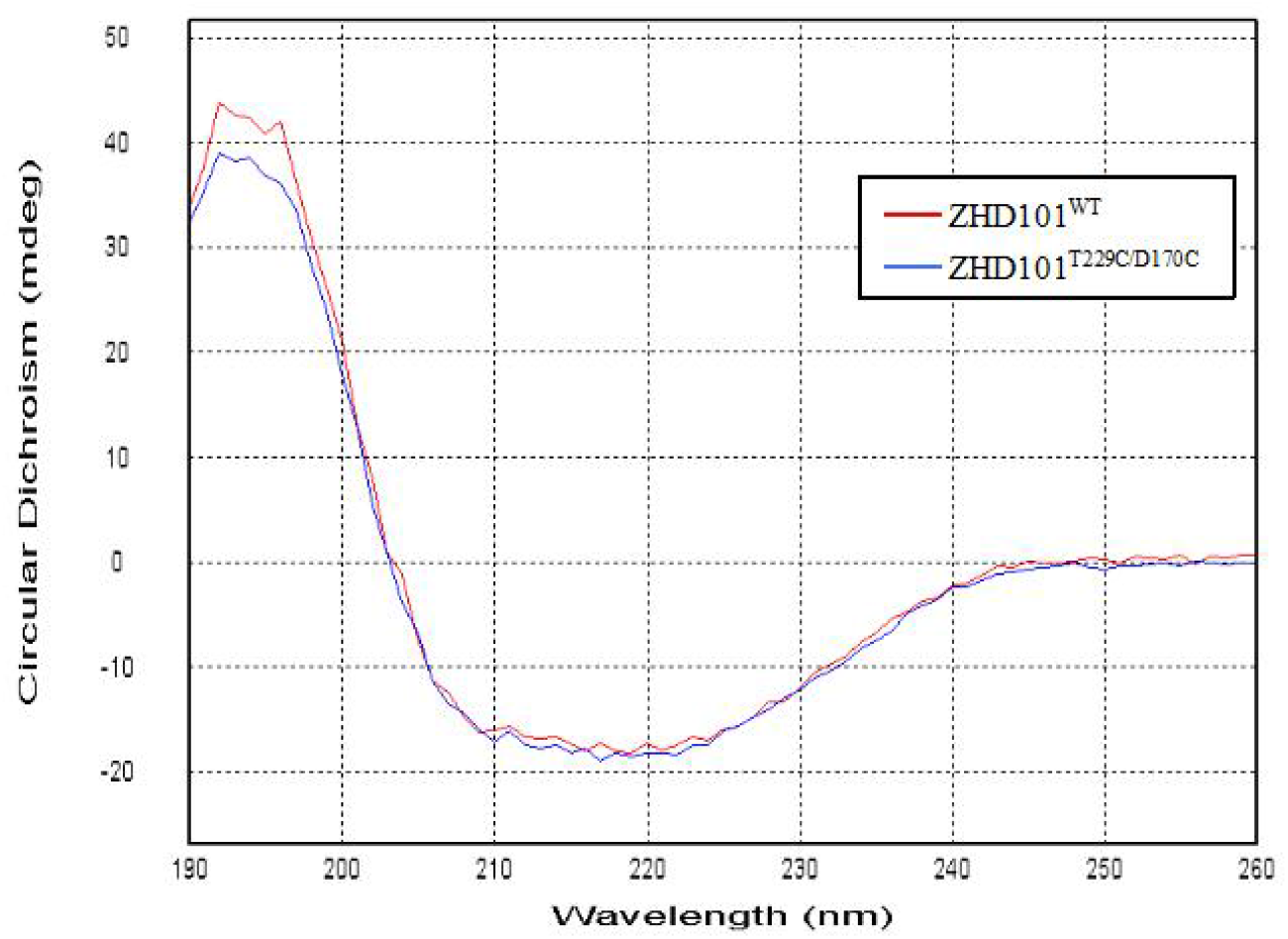
Full wavelength circular dichroism spectra of WT and ZHD101^T229C/D170C^. ZHD101^T229C/D170C^ with adjacent amino acids in two dimensions,(c) 170th amino acid of WT with adjacent amino acids, (d) 170th amino acid of ZHD101^T229C/D170C^ with adjacent amino acids.

**TABLE 1.**
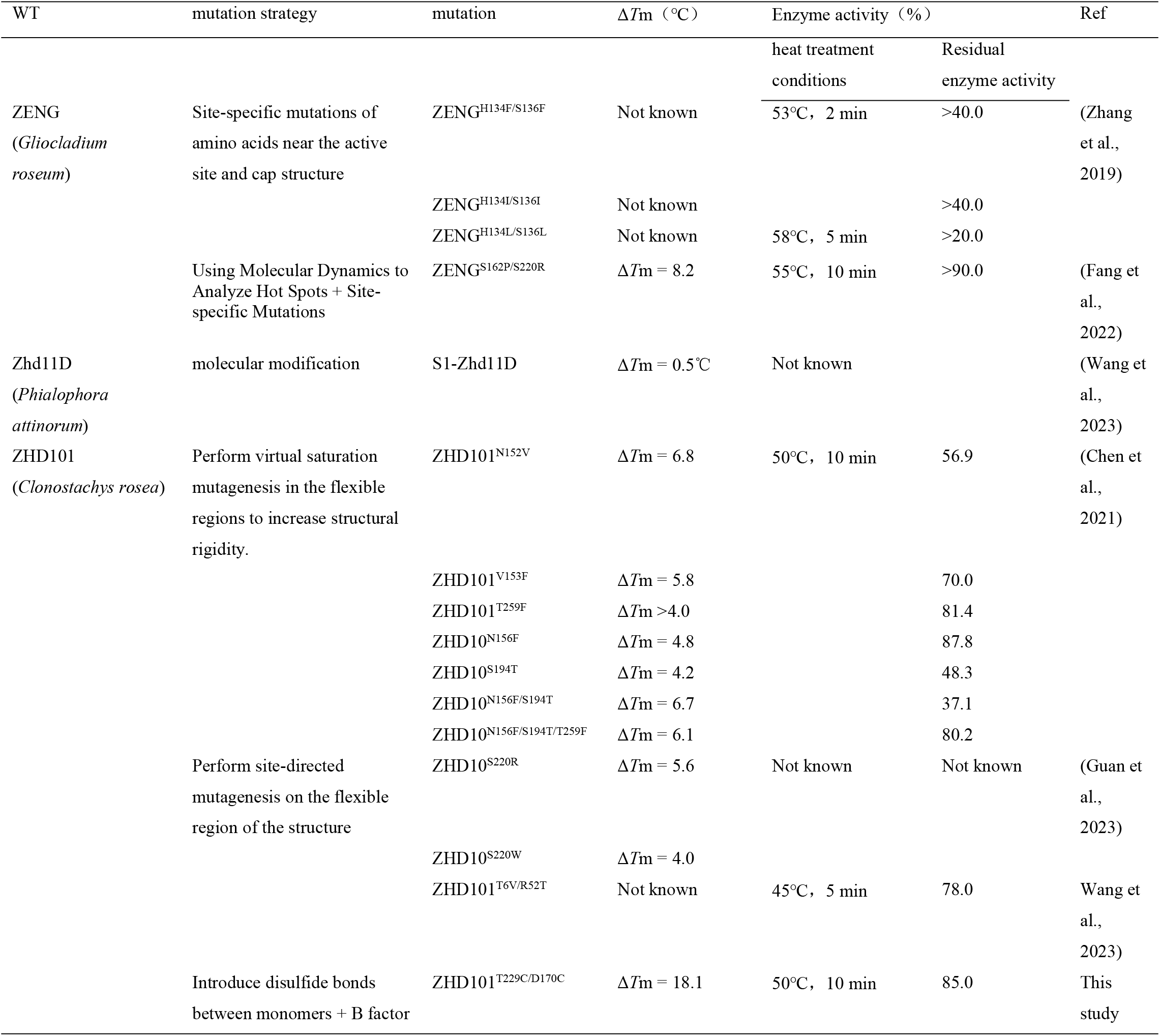
Summary of recent research on improving the thermal stability of ZEN lactonase.

Compared with other literature reports, it has the better improvement effect and is currently the highest value (*ΔT_m_*) in relevant research reports.

### Characteristic analysis of WT and ZHD101^T229C/D170C^

The optimal temperature, pH, and pH stability of the mutant enzymes were determined using the abovementioned methods. The optimal reaction temperature of ZHD101 and ZHD101^T229C/D170C^ was 40°C, and mutant enzyme activity was higher than that of the wild type at different temperatures (Fig S5(a)). The optimal reaction pH of ZHD101 and ZHD101^T229C/D170C^ was 8.5 and 7.5, respectively (Fig S5(b)). ZHD101 and ZHD101^T229C/D170C^ exhibited similar pH stability (Fig S5(c)). The kinetic parameters of the enzymes are listed in Table 2; there is no significant difference between WT and ZHD101^T229C/D170C^.

**TABLE 2.**
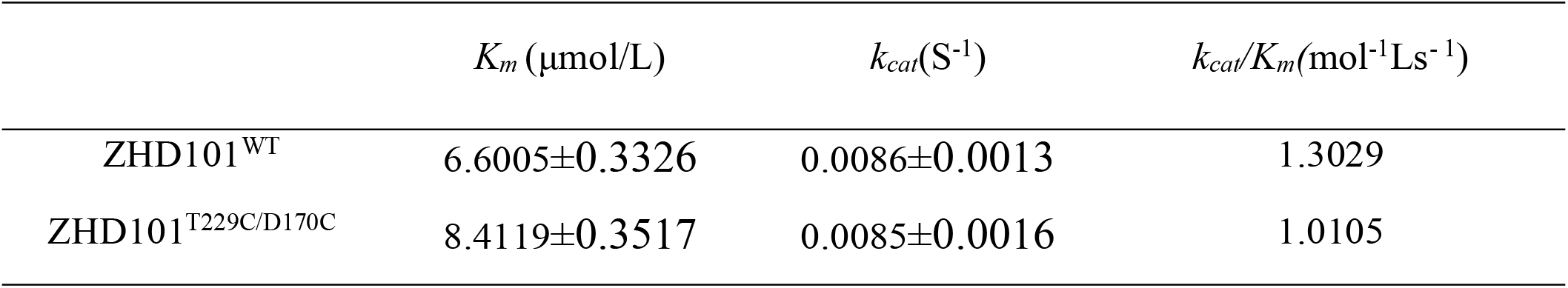
Kinetic parameters of wild-type and mutant ZHD101^T229C/D170C^.

### Interaction analysis of mutation sites

To further understand the reasons for the improved thermal stability of the mutant, the bonding situation near the mutation site was analyzed based on the structures of WT and ZHD101^T229C/D170C^. In the WT molecule, two ZHD101 monomers, A and B, aggregated into dimers through non-covalent interactions to exert their functions. Mutation of threonine to cysteine at the 229th position of the monomer could promote the formation of stable chemical bonds between monomers A and B, thereby improving the rigidity of the dimer. The 229 cysteine residues of the two monomers condensed to form a disulfide bond, consistent with our original design goal. Additionally, as shown in Fig 5(a-b), via alkyl interactions, cysteine at position 229 of monomer A could interact with lysine at position 230 and valine at position 226 of monomer B. These newly identified interactions increased the rigidity of the quaternary structure of the dimer. However, this mutation also caused a shift in the amino acid residues in this region, shortening the bonding distance between the amino acids in this region (Table 3).

**Figure 5.**
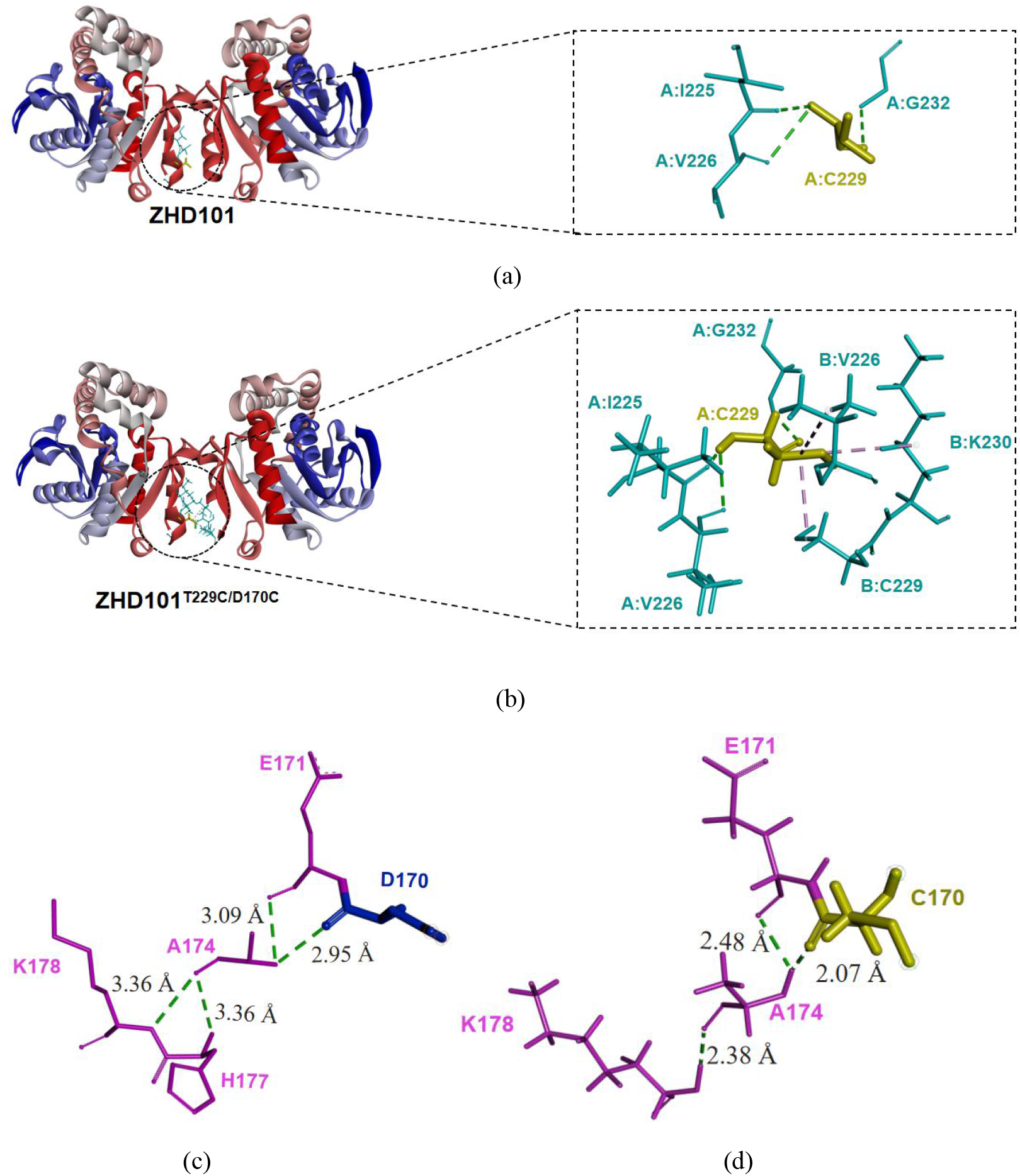
Bonding status of the (a) 229th amino acid of the wild type (WT) with adjacent amino acids in two dimensions, (b) 229th amino acid of ZHD101^T229C/D170C^ with adjacent amino acids in two dimensions,(c) 170th amino acid of WT with adjacent amino acids, (d) 170th amino acid of ZHD101^T229C/D170C^ with adjacent amino acids.

**TABLE 3.** Bonding situation of amino acids near the mutation site.

| Mutation site | Amino acids | Bonding type | Bond distance [Å] |  |
| --- | --- | --- | --- | --- |
|  |  |  | Wild type | ZHD101<sup>T229C/D170C</sup> |
| T229C | A: C229-A: I225 | Hydrogen bond | 3.00 | 1.99 |
|  | A: C229-A: V226 | Hydrogen bond | 3.24 | 2.76 |
|  | A: C229-B: G232 | Hydrogen bond | 3.28 | 2.41 |
|  | A: C229-B: C229 | Disulfide bond | – | 3.56 |
|  | A:C 229-B: K230 | Alkyl interaction | – | 4.91 |
|  | A: C229-B: V226 | Alkyl interaction | – | 4.73 |
| D170C | A: C170-A: A174 | Hydrogen bond | 2.95 | 2.07 |
|  | A: A174-A: E171 | Hydrogen bond | 3.09 | 2.48 |
|  | A: A174-A: K178 | Hydrogen bond | 3.36 | 2.38 |
|  | A: A174-A: H177 | Hydrogen bond | 3.36 | – |

As shown in Fig 5(c-d), in the wild type and mutant, the amino acid residue at position 170 on the molecular surface only formed a hydrogen bond with alanine at position 174; it did not interact with other amino acid residues. After the mutation of aspartic acid at position 170 to cysteine, the bonding distance between amino acid 170 and alanine 174 changed from 2.95 Å to 2.07 Å. At the same time, the bonding distance between alanine 174 and histidine lysine 178 was indirectly shortened (Table 3); this may have contributed to the improvement in thermal stability.

## Conclusion

This study obtained a "small and precise" library containing 3 mutants by introducing multiple designs such as disulfide bonds between monomers, B-factor analysis, and molecular dynamics simulation. The thermal stability of these mutants was evaluated and it was found that the thermal stability of ZHD101^T229C/D170C^ was significantly improved compared with the wild type (ΔTm = 18.1℃), and the catalytic efficiency was not negatively affected. This mutant enzyme is expected to be a favorable competitor in industrial feed additives, such as low-cost detoxification ZEN. The improvement strategy and identification method of this study can provide a reference for the thermal stability modification and optimization of other enzymes and proteins.

## Materials and Methods

### Strains, plasmids, and reagents

*Escherichia coli* (*E. coli*) DH5α was used as the cloning host strain. Plasmids pET28a (+) and *E. coli* BL21(DE3) were used for gene expression analysis. ZHD101 (GenBank ID: ABO076037.1) was synthesized using optimized codons from Shanghai Generay Biotech Co., Ltd. (Shanghai, China). for heterologous expression. Luria–Bertani (LB) medium was prepared to culture *Escherichia coli* cells. Zearalenone was purchased from Romer Labs (Tulln, Austria) and dissolved in acetonitrile to prepare a standard stock solution (5 mg/mL). All other chemical reagents were of analytical grade.

### Selection of disulfide bond formation sites between subunits

The 3D structure of ZHD101 (PDB ID: 3wzl) was downloaded from the PDB Protein Data Bank (http://www.rcsb.org/) and that of ZEN was downloaded from PubChem (https://pubchem.ncbi.nlm.nih.gov/). We used Discovery Studio 2023 (BIOVIA, USA, DS2023) for the molecular docking analysis, which was performed using the CHARMm-based docking program CDOCKER, with ZHD101 as the receptor and ZEN as the substrate. After docking, all amino acid residues in the binding zone of 5 Å between ZHD101 and ZEN were analyzed via alanine scanning, and the key amino acids affecting the binding, which must be avoided when designing mutation sites, were identified. We used two methods to predict the disulfide bonds: Method 1: DS4.5 software; Method 2: DbD2 website (http://cptweb.cpt.wayne.edu/DbD2/).

### Prediction of mutation sites based on B-factor

The B-factor values of each amino acid in the protein were evaluated using the isotropic displacement of DS2023, and candidate mutation sites were selected. Next, we determined the mutation energy (stability) to perform virtual amino acid mutations at candidate mutation sites. Finally, we calculated the folding free energy of wild-type proteins and mutants with mutation energy less than -2 kcal/mol and evaluated the structural stability of the mutants. The DynaMut web server was used for the computations.

### Protein expression and purification

*E. coli* BL21(DE3) transformants were cultured for more than 12 h at 37°C, after which 1% of the culture was inoculated into 300 mL fresh LB. The cultures were grown at 37°C to a cell turbidity of 0.4–0.6 at 600 nm and then induced for 24 h at 20°C by adding 0.5 mM IPTG. The cells were collected by centrifugation at 4°C for 10 min at 8 000 r/m. Next, the cells were resuspended in Tris-HCl buffer (20 mmol/L, pH 7.4) and ultrasonically disrupted on ice for 30 min (ultrasonic, 5 s; pause, 10 s); the lysates were then centrifuged at 4°C for 10 min at a speed of 12 000 rpm to remove cell debris. The recombinant enzyme with a His-tag was purified on a Ni^2+^-NTA agarose affinity column (General Electric Company, USA). The purified enzyme was concentrated and changed to the solution (20 mM Tris-HCl buffer, pH 7.4) using an ultrafiltration centrifuge tube (Millipore Sigma, USA), 4°C, 4000 × g, 20 min. The purified enzyme containing the enzyme was analyzed using 10% sodium dodecyl sulfate-polyacrylamide gel electrophoresis (SDS-PAGE) and collected for further experiments. The protein concentrations were measured using the Bradford method [36].

### Identification of disulfide bonds between protein subunits

Protein samples were prepared by separately mixing with an equal volume of SDS-PAGE loading buffer containing β-mercaptoethanol and loading buffer without β-mercaptoethanol. The samples were denatured by heating at 95°C for 5 min, and the weight of the two samples was compared using SDS-PAGE.

### Enzymatic activity assay

The degradation rate of ZEN characterized the activity of ZHD. The ZEN degradation assays were conducted in 50 mmol/L Tris-HCl buffer (pH 7.4) containing 25 μg/mL ZEN and 0.02 mg/mL purified enzyme. The reaction was carried out at 37°C for 10 min and subsequently terminated by adding 200 μL of methanol. Three replicates were performed for each group. The residual solution was filtered through a 0.22-µm filter and subjected to an HPLC system (UltiMate3000; Thermo Scientific, Bohemia, NY, USA) equipped with an ultraviolet detector (detection wavelength: 274 nm), and the injection volume was 20 μL. The flow rate of the mobile phase (at 15 min, 100% methanol; otherwise, 65% acetomethanol and 35% ultrapure water) was 1 mL/min. Concentrations of ZEN were determined based on retention times and peak areas compared to ZEN standard stock solution. The peak area vs. ZEN concentration standard curve was plotted and was in very good correlation. One unit of enzymatic activity was defined as the amount of enzyme required to degrade 1 μg of ZEN per min under optimal conditions.

### Analysis of enzyme thermostability and structure

The temperature at which residual enzyme activity decreased to 50% of its initial activity was its thermal half-inactivation temperature (*T_50_*). For measurement of the *T_50_* of the enzyme, the purified enzyme was incubated at 20, 30, 40, 50, and 60°C in Tris-HCl buffer (pH 7.4) for 10 min. Subsequently, the residual enzymatic activity was measured under optimal parameters. The time required for the residual enzyme activity to decrease to 50% of its initial activity is defined as the enzyme half-life (*t_1/2_*), also referred to as the thermal half-inactivation time. To measure the *t_1/2_*of the enzyme, the purified enzyme was incubated at 50°C in Tris-HCl buffer (pH 7.4) for 0, 1-, 2-, 5-, and 10-min. Enzymatic activity was measured using the optimal parameters. All experiments were conducted in triplicate. Wild-type enzyme activity was 100%, and the relative enzyme activity of other groups was calculated.

The secondary structure of the enzyme was evaluated via CD spectroscopy, and the thermal melting temperature (*T_m_*) was determined using thermal denaturation experiments. The purified sample (0.1 mg/mL) was scanned at 25°C at full wavelength in the 190–260 nm range, with an increase of 0.5 nm every 2 ns. Characteristic absorption peaks measured across the full wavelength range were used to determine the secondary structure of the protein. Next, a thermal denaturation experiment was conducted, setting the temperature range from 20°C to 100°C at a heating rate of 2°C per min. Changes in the CD signal at 222 nm were measured for each enzyme. By plotting the CD signal against the temperature, the midpoint of this transition, where half of the protein unfolds, was determined to be the *T_m_*.

### Characterization of wild-type and mutant enzymes

In the optimal temperature assay, enzymatic activity was measured at 20, 30, 40, 50, and 60°C in Tris-HCl buffer (pH 7.4). In the optimal pH assay, the enzyme was assayed in different buffers, including 20 mM phosphate buffer (pH 4.5, 5.5, and 6.5), 20 mM Tris-HCl buffer (pH 7.5 and 8.5), and 20 mM carbonate buffer solution (pH 9.5 and 10.5) at 37°C. For pH stability, the enzyme was pre-incubated in different buffers, including 20 mM phosphate buffer (pH 4.5, 5.5, and 6.5), 20 mM Tris-HCl buffer (pH 7.5 and 8.5), and 20 mM carbonate buffer solution (pH 9.5 and 10.5) for 30 min, and enzyme activity was measured under optimal parameters. The initial reaction rate was calculated from the slope of the zero-order plot of product concentration versus reaction time, and the initial reaction rate was then plotted against the substrate concentration. The resulting curve was fitted to the hyperbolic equation v =*V_max_*×[*S*]/(*K_m_*+[*S*]) using Origin 7.0 (OriginLab, Northampton, MA, USA) to obtain the turnover number (*k_cat_*) and *K_m_*. All measurements were duplicated with substrate concentrations ranging from 10 to 500 mM.

## ACKNOWLEDGMENTS

This study was supported by the National Key Research and Development Program of China (grant number 2021YFC2103003). W. D. drafted the manuscript and analyzed the data. Y.H. performed the enzymatic experiments. H.Z. performed molecular docking analysis under the supervision of C.X. S.Z. and D.S. designed the experiments and supervised the project.

## Data availability

Data will be made available on request.

## FUNDING

Funder Grant(s) Author(s)

National Key Research and Development Program of China 2021YFC2103003 Dongsheng Yao

## AUTHOR CONTRIBUTION

Weiqiu Ding, Writing – original draft, Methodology, Data curation, Conceptualization. Yujie Huang, Methodology, Conceptualization. Haiyi Zhang, Software, Formal analysis. Shaoyan Zheng, Writing – review & editing. Chunfang Xie, Formal analysis, Data curation. Dongsheng Yao, Supervision, Funding acquisition.

